# Nucleus accumbens dopamine encodes the trace period during appetitive Pavlovian conditioning

**DOI:** 10.1101/2025.01.07.631806

**Authors:** Matthew J. Wanat, Brandon I. Garcia-Castañeda, Cecilia Alducin-Martinez, Leonor G. Cedillo, Erika T. Camacho, Paul E.M. Phillips

**Author notes:** **Corresponding Author:** Matthew J. Wanat, Department of Neuroscience, Developmental and Regenerative Biology University of Texas at San Antonio. Authors contributed equally to the work (Brandon I. Garcia-Castañeda, Cecilia Alducin-Martinez, Leonor G. Cedillo). **Author Contributions:** MJW and PEMP designed the experiments. MJW, BIG, CA, and LGC collected the data. MJW, BIG, CA, and ETC analyzed the data. All authors assisted with writing the manuscript.

## Abstract

Pavlovian conditioning tasks have been used to identify the neural systems involved with learning cue-outcome relationships. In delay conditioning, the conditioned stimulus (CS) overlaps or co-terminates with the delivery of the unconditioned stimulus (US). Prior studies demonstrate that dopamine in the nucleus accumbens (NAc) regulates behavioral responding during delay conditioning. Furthermore, the dopamine response to the CS reflects the relative value of the upcoming reward in these tasks. In contrast to delay conditioning, trace conditioning involves a ‘trace’ period separating the end of the CS and the US delivery. While dopamine has been implicated in trace conditioning, no studies have examined how NAc dopamine responds to reward-related stimuli in these tasks. Here, we developed a within-subject trace conditioning task where distinct CSs signaled either a short trace period (5s) or a long trace period (55s) prior to food reward delivery. Male rats exhibited greater conditioned responding and a faster response latency to the Short Trace CS relative to the Long Trace CS. Voltammetry recordings in the NAc found that the CS-evoked dopamine response increased on Short Trace trials but decreased on Long Trace trials. Conversely, US-evoked dopamine responses were greater on Long Trace trials relative to Short Trace trials. The CS dopamine response correlated with the response latency and not with conditioned responding. Furthermore, the relationship between CS dopamine and latency was best explained by an exponential function. Our results collectively illustrate that the trace period is encoded by the bidirectional NAc dopamine response to the CS during Pavlovian conditioning.

**Significance statement:** Learning to associate a cue with given outcome is a fundamental process underlying reward seeking behavior. Striatal dopamine is important for associating cues with rewards during Pavlovian conditioning. However, it is unclear how the dopamine system responds to cues during trace conditioning when there is temporal gap between the cue and reward. Here, we performed voltammetry recordings of striatal dopamine levels in male rats during trace conditioning. We find that cue-evoked dopamine signals encode the trace period and is related to the response latency. While prior reports find dopamine neurons signal the relative reward value by increases in dopamine levels, we demonstrate that the dopamine response to reward-predictive cues can signal the reward value through bidirectional changes in dopamine transmission.

## Introduction

Associating a cue with a given outcome is a fundamental process critical for both reward learning and aversive learning. These learning processes are commonly studied in the context of Pavlovian conditioning tasks in which a conditioned stimulus (CS) predicts the delivery of an unconditioned stimulus (US). Delay conditioning is a category of Pavlovian conditioning in which the presentation of the CS overlaps or co-terminates with the delivery of the US. Pharmacological and optogenetic studies illustrate that dopamine signaling facilitates learning and the expression of conditioned responses during delay conditioning tasks with food rewards (Di Ciano et al., 2001; Flagel et al., 2011; Saunders and Robinson, 2012; Lopez et al., 2015; Fraser et al., 2016; Fraser and Janak, 2017; Roughley and Killcross, 2019; Heymann et al., 2020; Lee et al., 2020; Morrens et al., 2020; Stelly et al., 2020; Khoo et al., 2021; Stelly et al., 2021; Lefner et al., 2022b). Recordings from midbrain dopamine neurons as well as dopamine release in the nucleus accumbens (NAc) demonstrate that the CS-evoked dopamine response signals the relative value of the upcoming reward in delay conditioning tasks (Fiorillo et al., 2003; Tobler et al., 2005; Hart et al., 2015; Fonzi et al., 2017; Lefner et al., 2022a). Furthermore, the direction of the CS-evoked dopamine response signals the valence of the outcome, as NAc dopamine levels decrease in response to the CS in delay conditioning tasks with an aversive US (Matsumoto and Hikosaka, 2009; Badrinarayan et al., 2012; Oleson and Cheer, 2013; de Jong et al., 2019; Stelly et al., 2019; Kutlu et al., 2021; Salinas-Hernandez et al., 2023; Lopez and Lerner, 2025).

In contrast to delay conditioning where there is no temporal separation between the CS and US, trace conditioning tasks involve a ‘trace’ period following the end of the CS and the delivery of the US. While many trace conditioning tasks commonly use an aversive stimulus for the US (e.g. eyeblink conditioning, conditioned taste aversion, and fear conditioning), trace conditioning tasks can also use food rewards for the US (Reilly and Bornovalova, 2005; Woodruff-Pak and Disterhoft, 2008; Cohen et al., 2012; Raybuck and Lattal, 2014; Pezze et al., 2015; Eshel et al., 2016; Sias et al., 2024). Pharmacological and genetic studies indicate that dopamine signaling regulates behavioral responding during appetitive and aversive trace conditioning tasks (Fenu et al., 2001; Fenu and Di Chiara, 2003; Deng et al., 2015; Pezze et al., 2015). However, despite the evidence for dopamine’s involvement, no studies to date have examined how NAc dopamine signals relate to behavioral outcomes during appetitive trace conditioning.

In this study, male rats were trained on a trace conditioning task where distinct audio CSs signaled either a Short Trace period (5s) or a Long Trace period (55s) prior to the delivery of a food reward. Voltammetry recordings of NAc dopamine release were performed in well-trained animals. This experimental approach allows for the within-subject analysis of NAc dopamine signals to two distinct cue-outcome pairs during trace conditioning. If CS-evoked dopamine response reflects reward value, one would anticipate a larger dopamine response to the Short Trace CS (greater value for an imminent reward) relative to the Long Trace CS (lower value for a delayed reward). Consistent with this prediction, we observed a larger CS-evoked dopamine response on Short Trace trials, though notably there was a decrease in dopamine levels on Long Trace trials. Subsequent analyses found that CS-evoked changes in dopamine transmission were related to response latency according to an exponential decay function. Overall, these data illustrate notable differences in the CS-evoked NAc dopamine response between delay and trace conditioning tasks. In delay conditioning tasks, the increase in CS-evoked dopamine response scales with reward value (Fiorillo et al., 2003; Tobler et al., 2005; Fonzi et al., 2017; Lefner et al., 2022a). In contrast, we find that NAc dopamine levels decrease in response to a reward-predicting CS associated with a Long Trace period. Therefore, dopamine encoding of reward value is not solely limited to positive changes in dopamine levels, but rather can encompass both positive and negative changes in the dopamine response to the CS.

## Methods

### Subjects and surgery

All procedures were approved by the Institutional Animal Care and Use Committee at the University of Washington and the University of Texas at San Antonio. For voltammetry recording experiments, male Sprague-Dawley rats (Charles River) aged P60-65 were pair-housed upon arrival and given ad libitum access to water and chow and maintained on a 12-hour light/dark cycle (n = 23). We acknowledge the limitation of only including male subjects. Though we note that the in vivo dopamine recordings for this study were collected in 2011-2013, which was prior to the NIH policy regarding sex as a biological variable, when it was regrettably commonplace to perform single sex studies. Rats underwent surgery to implant voltammetry electrodes in the NAc core (relative to bregma: 1.3 mm anterior; ± 1.3 mm lateral; 7.0 mm ventral) and a Ag/AgCl reference electrode. Carbon fiber voltammetry electrodes consisted of a carbon fiber housed in silica tubing and cut to a length of ~150 µm (Clark et al., 2010). Rats were single-housed following surgery. Additional behavior-only experiments were performed using male and female Sprague-Dawley rats (Inotiv).

### Behavioral procedures

All procedures were performed during the light cycle. After >1 week of recovery from surgery, rats were placed and maintained on mild food restriction (~15 g/day of standard lab chow) to target 90% free-feeding weight, allowing for an increase of 1.5% per week. Behavioral sessions were performed in chambers (Med Associates) that had a house light, a food tray, and auditory stimulus generators (whitenoise and 2.9 kHz tone). To familiarize rats with the chamber and food retrieval, rats underwent a single magazine training session in which 20 food pellets (45 mg, BioServ) were non-contingently delivered at a 90 ± 15 s variable interval. Trace conditioning sessions consisted 25 Short Trace trials and 25 Long Trace trials that were presented in a pseudorandom manner. Short Trace trials consisted of a 5 s audio CS followed by a 5 s trace period after which a single food pellet was delivered. Long Trace trials consisted of a 5 s audio CS followed by a 55s trace period after which a single food pellet was delivered. The food tray light illuminated for 4.5 s after the delivery of the food pellet US and trials were separated by a 5 -15 s intertrial interval (ITI). The audio CSs used for Short and Long Trace trials (tone and whitenoise) were counterbalanced across subjects. Rats underwent a total of 30 trace conditioning sessions. Rats trained on an unrewarded version of the trace conditioning task underwent the identical procedures except that rewards were not delivered.

This study included a reanalysis of published data from rats trained on a related delay conditioning task (Fonzi et al., 2017). Male rats from this previous study underwent surgery and were placed under food restriction as outlined above. Delay conditioning sessions consisted of 50 trials where the termination of the 5 s audio CS (whitenoise or 2.9 kHz tone, counterbalanced across animals) resulted in the delivery of a single food pellet and illumination of the food tray light for 4.5 s. The distinct CSs in this delay conditioning task signaled the duration of the preceding ITI, with rats receiving 25 Short Wait CS trials and 25 Long Wait CS trials presented in a pseudorandom manner. Our reanalysis focused on a subset of Short Wait CS trials that had an equivalent ITI duration to the rats that were trained on the trace conditioning task (e.g. 10s and 15s ITI). Behavioral metrics monitored include the head entry latency following the onset of the CS presentation, the head entry latency following the US delivery, and conditioned responding. We calculated conditioned responding as the rate of head entries during the 5 s CS relative to the rate of head entries during the 5 s preceding the CS, consistent with prior studies (Fonzi et al., 2017; Stelly et al., 2020; Stelly et al., 2021; Lefner et al., 2022a; Lefner et al., 2022b).

### Voltammetry recordings

Voltammetry recordings of NAc dopamine release were performed in well-trained subjects (sessions 25-30). Chronically-implanted electrodes were connected to a head-mounted amplifier to monitor changes in dopamine release in behaving rats using fast-scan cyclic voltammetry as described previously (Clark et al., 2010; Fonzi et al., 2017; Oliva and Wanat, 2019; Stelly et al., 2019; Stelly et al., 2020; Oliva et al., 2021; Stelly et al., 2021; Lefner et al., 2022a). The carbon fiber electrodes were held at -0.4 V (vs. Ag/AgCl) with voltammetric scans applied at 10 Hz in which the potential was ramped in a triangular waveform to +1.3 V and back to -0.4 V at a rate of 400 V/s. A principal component regression analysis was performed on the voltammetry signal using a standard training set (Heien et al., 2005). The average post-implantation sensitivity of electrodes (34 nA/µM) was used to estimate dopamine concentration (Clark et al., 2010). Chemical verification of dopamine was achieved by obtaining a high correlation of the cyclic voltammogram during a reward-related event to that of a dopamine standard (correlation coefficient r^2^ ≥ 0.75 by linear regression). Voltammetry data for a session were excluded from analysis if the detected voltammetry signal did not satisfy this chemical verification criteria (Fonzi et al., 2017; Oliva and Wanat, 2019; Stelly et al., 2019; Stelly et al., 2020; Oliva et al., 2021; Stelly et al., 2021; Lefner et al., 2022a). Voltammetry data from given electrode was included provided there were two or more recording sessions that satisfied the above stated inclusion criteria (n = 10 electrodes). The CS-evoked dopamine response was quantified as the average dopamine signal during the 5 s CS relative to the 5 s prior to the CS. The US-evoked dopamine response was quantified as the average dopamine signal during the 3 s following the US delivery relative to the average dopamine signal during the 1 s preceding the US delivery. The reported dopamine signals therefore reflect the relative change in dopamine levels to a baseline period immediately preceding the event of interest.

### Modeling and data analysis

We examined the relationship between the CS dopamine response and latency as a function of the trace period. Data from the full trace conditioning data set were fit to a linear function with 2 parameters:

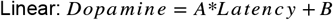

Prior research indicates that exponential and hyperbolic functions can account for the relationship between cue-evoked dopamine neuron firing and reward delay in primates (Kobayashi and Schultz, 2008). To perform non-linear curve fits on the current data set, we reexamined data collected from a related delay conditioning task where the trace period was 0 s (Fonzi et al., 2017). This delay conditioning task used the same audio CSs and delivered the same number of rewards per session as the trace conditioning task. Delay conditioning rats also underwent the same training procedure with voltammetry recordings performed only in well-trained animals (>24 conditioning sessions). We solely focused on trials with the same ITI duration in both the delay and trace conditioning tasks (10 and 15 s ITI). In this manner we examined how the trace period (0, 5, 55 s) influences the CS dopamine response and its relationship to response latency. We note that hyperbolic functions are not suitable for our data set which includes negative values for the CS dopamine response on Long Trace trials. We therefore initially fit the data from trace and delay conditioning rats to an exponential function with 3 parameters:

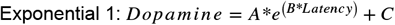

This Exponential 1 function gave a better fit and smaller error than single exponential function with 2 parameters. In all cases the parameters were optimized to give the best fit without restriction on the parameters. When examining the individual data for each trace period (0, 5, 55 s) utilizing the Exponential 1 function, we found (i) the fit for the delay conditioning / 0 s trace data to be concave up and increasing, (ii) the fit for the 5 s trace data to be concave down and decreasing, and (iii) the fit for the 55 s trace data to be concave up and decreasing. Taken together, this dopamine-latency relationship suggested a combined behavior similar to a Gaussian function. Thus, the data were also fit to an exponential function with 4 parameters (specifically, a Gaussian function) that was optimized to account for the dopamine-latency relationship within each trace period (0, 5, 55 s):

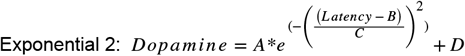

The performance of these functions was evaluated on the full data set from the trace conditioning rats. The deviation of the predicted dopamine versus actual dopamine values were quantified for each equation.

Statistical analyses were performed in GraphPad Prism. Significant effects were determined by performing repeated measures ANOVAs with post-hoc Sidak’s tests where appropriate, paired t-tests, and linear regressions. The Geisser-Greenhouse correction was applied to address unequal variances between groups. A Friedman test followed by a post-hoc Dunn’s test was used to examine data that did not conform to a Gaussian distribution.

## Results

Rats were trained on a trace conditioning task that contained two distinct trial types (**Fig. 1A**). During Short Trace trials, the presentation of a 5 s audio CS (tone or whitenoise) was followed by a 5 s trace period after which the US was delivered (food pellet). During Long Trace trials, the presentation of a 5 s audio CS (whitenoise or tone) was followed by a 55 s trace period after which the US was delivered. The assignment of the audio CS to a given trial type was counterbalanced between animals. Sessions consisted of a 25 Short Trace trials and 25 Long Trace trials that were presented in a pseudorandom pattern and separated by a 5 - 15 s ITI. Rats underwent 24 behavior-only training sessions to ensure they were well-trained prior to performing voltammetry recordings. Animals exhibited higher levels of conditioned responding on Short Trace trials relative to Long Trace trials (two-way repeated measures ANOVA, effect of trial type F_(1,22)_ = 6.5, p = 0.018; interaction effect F_(4.4, 97.4)_ = 3.0, p = 0.018, n = 23; **Fig. 1B**). Importantly, the increase in conditioned responding on Short Trace trials reflects a learned association, as there was no increase in cue-driven behavior in rats trained on an unrewarded variant of the task (**Extended Data Fig. 1-1**). Despite the low level of conditioned responding on Long Trace trials, there was an increase in the head entries throughout the trace period that emerged over training (**Extended Data Fig. 1-2**). These data are consistent with a behavioral response that the rats have learned the temporal dynamics of the task and are anticipating the upcoming reward. The latency to initiate a head entry following the CS onset was longer on Long Trace trials relative to Short Trace trials (two-way repeated measures ANOVA; effect of trial type F_(1,22)_ = 26.4, *** p < 0.0001; interaction effect F_(6.0, 132.1)_ = 2.6, p = 0.021; **Fig. 1C**). Interestingly, this effect was driven by a slowing of the latency to respond on Long Trace trials. Finally, we examined the latency to the first head entry following the US delivery and found no difference between trial types (two-way repeated measures ANOVA; effect of trial type F_(1,22)_ = 1.0, p = 0.32; **Fig. 1D**). Collectively, these data illustrate that the presentation of the Short Trace CS enhances conditioned responding, whereas the presentation of the Long Trace CS delays the latency to perform a head entry.

**Figure 1.**
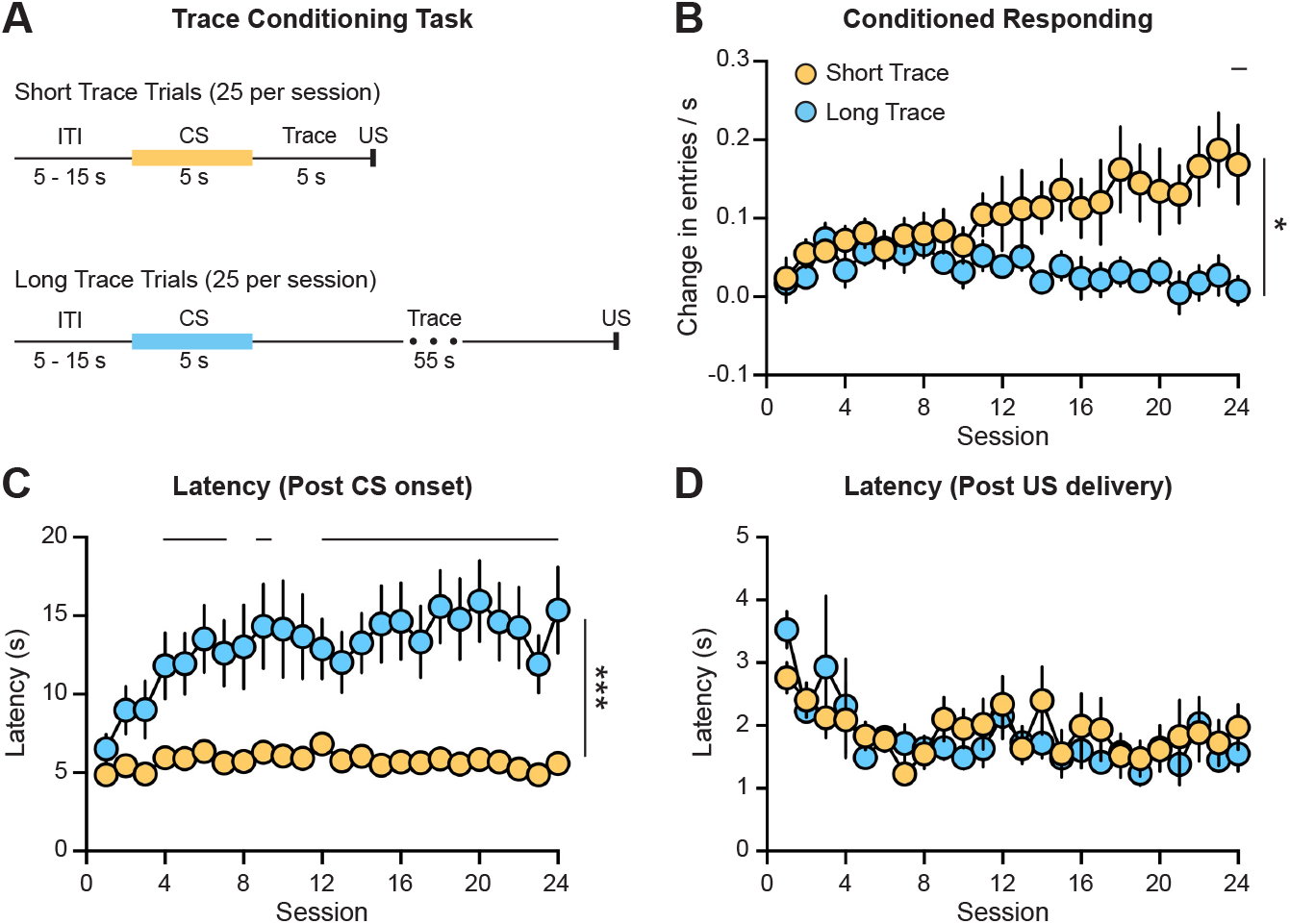
Trace conditioning task. (A) Schematic of Short and Long Trace trials, which are presented in a pseudorandom pattern within a session. (B) Higher levels of conditioned responding on Short Trace trials relative to Long Trace trials (effect of trial type, * p = 0.018). (C) Longer latency to perform a head entry following the CS onset on Long Trace trials relative to Short Trace trials (effect of trial type, *** p < 0.0001). (D) No difference in the latency to retrieve the reward between Short and Long Trace trials. Horizontal lines on graphs denote post hoc Sidak’s test, p < 0.05.

Voltammetry recordings of NAc dopamine levels were performed in rats that were well-trained on the trace conditioning task (>24 sessions). Data were analyzed from animals with electrodes in the NAc core with at least 2 viable voltammetry recording sessions (n = 15 electrodes from 10 rats; **Fig. 2A**). Dopamine levels increased in response to the Short Trace CS but decreased to the Long Trace CS (**Fig. 2B-C**). We quantified CS-evoked dopamine response as the average dopamine signal during the 5 s CS relative to the 5 s prior to the CS presentation. Dopamine release to the Short Trace CS was greater than the dopamine response to the Long Trace CS (paired t-test, t_14_ = 4.0, ** p = 0.001, **Fig. 2E**). Therefore, the magnitude of the CS-evoked dopamine response is inversely related to the trace period before the US is delivered. We also examined the CS dopamine response along with the first 5 s of the trace period and found a similar relationship with greater dopamine levels on the Short Trace trials relative to Long Trace trials (paired t-test, t_14_ = 3.1, ** p = 0.008, **Fig. 2F**). The dopamine response during this initial 5 s of the trace period on Long Trace trials remained stable (**Extended Data Fig. 2-1**), which suggests the decrease in dopamine levels is driven by the conditioned properties of the CS and not by a nonspecific drift in the dopamine signal. In contrast to the CS-evoked dopamine signals, there was a larger dopamine response to the US delivery on Long Trace trials relative to Short Trace trials (paired t-test, t_14_ = 3.4, ** p = 0.004). These findings are consistent with prior studies illustrating a greater dopamine response to delayed or unexpected rewards (Fiorillo et al., 2003; Wanat et al., 2010). In support, the magnitude of the US-evoked dopamine response on Long Trace trials was significantly smaller than the dopamine response to an unexpected reward delivered outside of the Pavlovian conditioning session (**Extended Data Fig. 2-2**).

**Figure 2.**
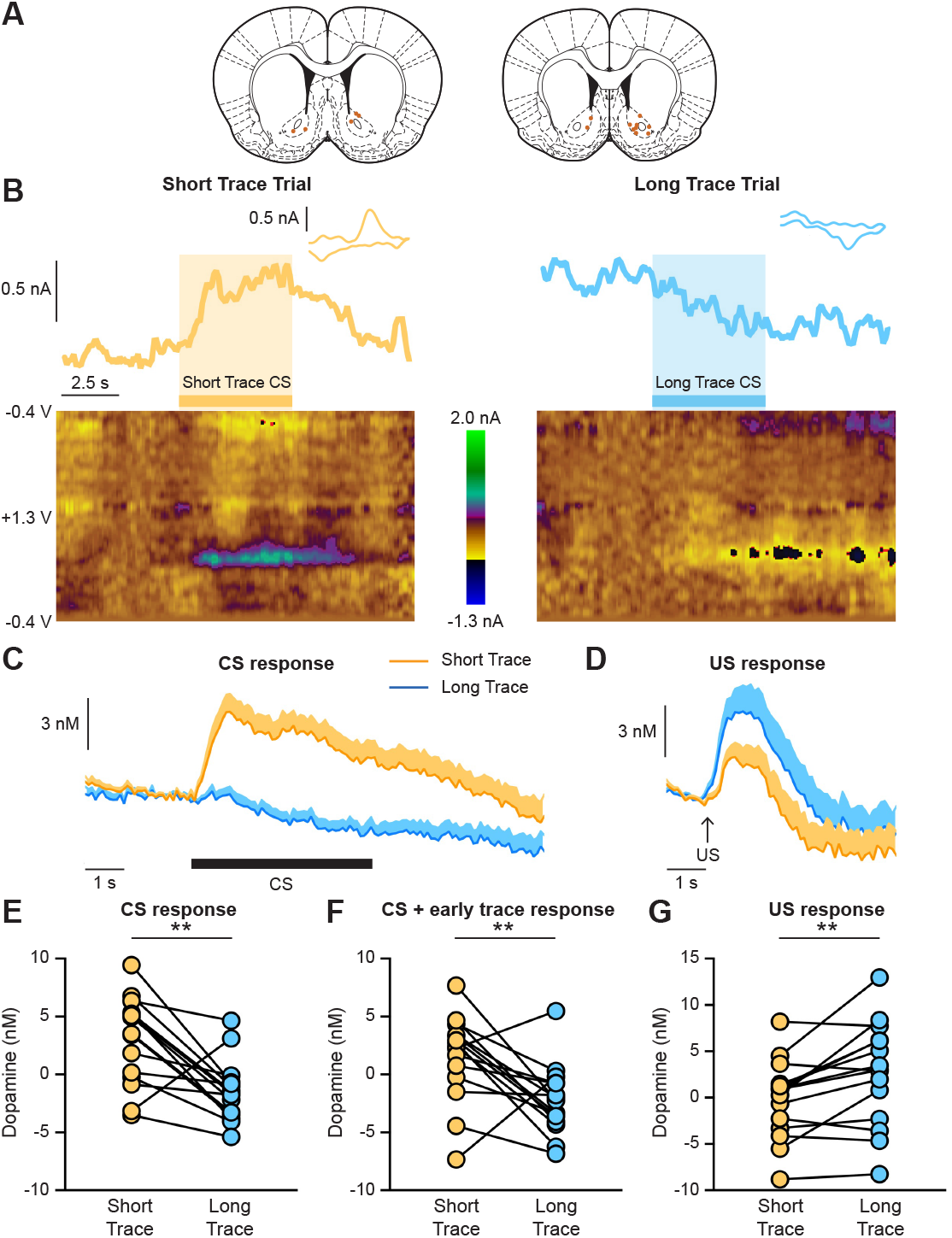
NAc dopamine release during trace conditioning. (A) Voltammery electrode placement. Voltammetry recordings were performed in well-trained rats (> 24 sessions) (B) Representative voltammetry recordings illustrating the dopamine current (top), cyclic voltammogram (top, inset), and the voltammetry color plot (bottom) from a Short Trace and Long Trace trial. (C) Average dopamine response to the CS and the initial trace period. (D) Average dopamine response to the US delivery. (E) Larger dopamine response to the Short Trace CS relative to the Long Trace CS (paired t-test, t_14_ = 4.0, ** p = 0.001). Dopamine to the Short Trace CS was greater than null values (one sample t-test relative to 0, t_14_ = 3.0, ** p = 0.009). (F) Larger dopamine response to the CS and the first 5 s of the trace period on Short Trace trials relative to Long Trace trials (paired t-test, t_14_ = 3.1, ** p = 0.008). Dopamine on Long Trace trials decreased below null values (one sample t-test relative to 0, t_14_ = 2.9, p = 0.01). (G) Larger dopamine response to the US delivery on Long Trace trials relative to Short Trace trials (paired t-test, t_14_ = 3.4, ** p = 0.004).

We next determined how the dopamine signals correlated with behavioral outcomes. Although Short Trace trials produced higher levels of conditioned responding and larger CS-evoked dopamine release relative to Long Trace trials (**Figs. 1-2**), there was no correlation between CS dopamine levels and conditioned responding (**Fig. 3A, left**). We also calculated the slope of this relationship for each electrode and found that the average slope was not significantly different from 0 (**Fig. 3A, right**). These data collectively illustrate the lack of a relationship between conditioned responding and CS dopamine release during trace conditioning. In contrast, we found that CS dopamine levels were inversely related to the head entry latency (r = -0.51, p = 0.004) and the slope of this relationship was significantly different from 0 across electrodes (one sample t-test relative to 0, t_14_ = 3.3, ** p = 0.005; **Fig. 3B**). Similar results were observed when examining the relationship between head entry latency and the dopamine response during the CS and the first 5 s of the trace period (r = -0.51, p = 0.004; **Fig. 3C**). While there was a larger dopamine response to the US on Long Trace trials, this was not related to the latency to retrieve the food reward (**Fig. 3D**).

**Figure 3.**
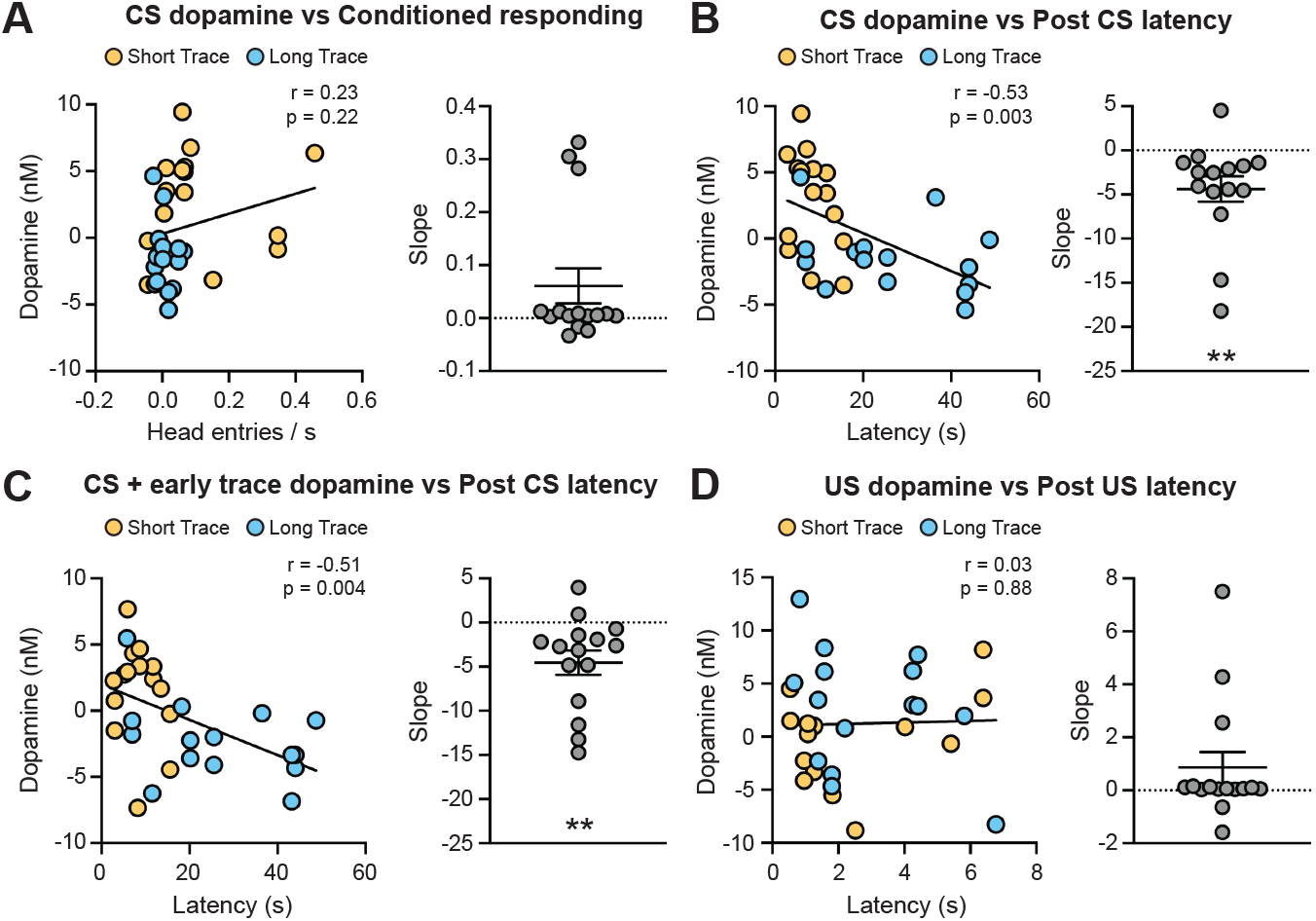
Correlating NAc dopamine signals to behavioral responding. Subpanels present the correlation between dopamine and behavior (left) and the slope of the relationship across electrodes (right). (A) The CS dopamine response was not correlated with conditioned responding. (B) The CS dopamine response was inversely related to the head entry latency (r = -0.53, p = 0.003). The slope of this relationship was significant across electrodes (one sample t-test relative to 0, t_14_ = 3.1, ** p = 0.009). (C) The dopamine response to the CS and the first 5 s of the trace period was inversely related to the head entry latency (r = -0.51, p = 0.004). The slope of this relationship was significant across electrodes (one sample t-test relative to 0, t_14_ = 3.3, ** p = 0.005). (D) The US dopamine response was not correlated with latency to retrieve the reward.

Prior studies indicate that linear and non-linear functions can account for the relationship between dopamine transmission and behavioral responding in timing-related tasks. The subjective hazard rate, or rather the expectation than an event will occur, is inversely related behavioral responding and dopamine neuron firing activity according to a linear function over rapid timescales (< 2 s) (Pasquereau and Turner, 2015). However, recordings from dopamine neurons during a delay discounting task using longer timescales (1 – 16s) demonstrated that the CS dopamine response decreases in a non-linear manner as the reward delay increases (Kobayashi and Schultz, 2008). Specifically, the decay in the CS dopamine response is steep for more immediate rewards and shallow for more delayed rewards (Kobayashi and Schultz, 2008). While there was a significant correlation between CS dopamine levels and head entry latency (**Fig. 3B**), the nature of this relationship may not be linear given the longer timescales used in the trace conditioning task. Determining how the trace period influences the CS dopamine response and its relationship to behavior therefore requires additional data from short trace periods when the decay in the dopamine response is steepest. To address this, we reexamined data collected from a related delay conditioning task where the trace period is 0 s (Fonzi et al., 2017). This delay conditioning task used the same audio CSs and delivered the same number of rewards per session as the trace conditioning task. Delay conditioning rats also underwent the same training procedure with voltammetry recordings performed only in well-trained animals (>24 conditioning sessions). We solely focused on trials with the same ITI duration in both the delay and trace conditioning tasks (10 and 15 s ITI). In this manner we can establish how the trace period (0, 5, 55 s) influences the CS dopamine response and its relationship to response latency (**Fig. 4A**).

**Figure 4.**
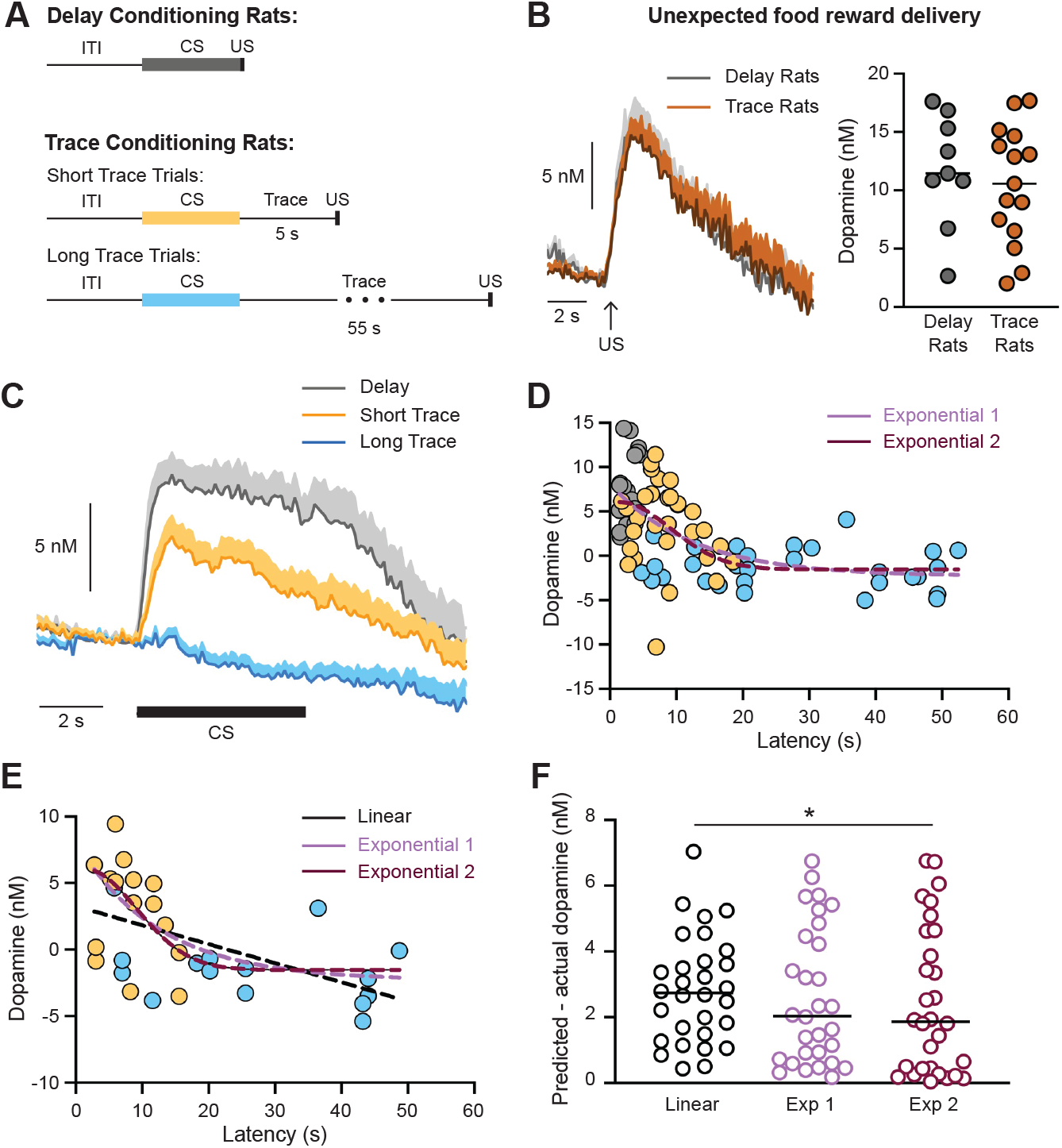
The relationship between CS dopamine and the latency to respond across different trace period durations. (A) Schematic of delay conditioning trials (with no trace period) and trace conditioning trials. (B) No difference in the dopamine response to unpredicted reward presentations between rats trained on the delay conditioning task or the trace conditioning task. (C) Average NAc dopamine release to the CS. (D) Data from 10 and 15 s ITI trials from rats trained on the delay and trace conditioning tasks. Overlayed dashed lines illustrate the the best fit exponential functions based on the delay and trace conditioning data using three variables (Exponential 1: *A* = 10.43, *B* = 0.07954, *C* = -2.32; pink) or four variables (Exponential 2: *A* = 7.568, *B* = -1.328, *C* = 11.06, *D* = -1.531; red). (E) Best fit exponential and linear functions overlayed on the average responses from the full data set (see also **Fig. 3B**; Linear function: *A* = -0.1429, *B* = 3.262; black). (F). Difference between the predicted and actual dopamine values for the linear and exponential functions (Friedman test, F = 6.07, p = 0.04; post-hoc Dunn’s test, * p = 0.04).

It is important to demonstrate that a given stimulus produces an equivalent dopamine response in both trace and delay conditioned rats before one compares the dopamine signals between these experiments. To address this, we evaluated the dopamine response to an unexpected food reward that was delivered outside of the training sessions. We found no difference in the NAc dopamine response to unexpected food rewards between trace and delay conditioned rats (**Fig. 4B**). As would be expected, CS-evoked dopamine levels in the NAc were greatest for delay conditioned rats with no trace period (**Fig. 4C**). We fit the data from 10 and 15s ITI trials to best-fit exponential functions with either three parameters (Exponential 1) or four parameters (Exponential 2; **Fig. 4D**). We then assessed the performance of these functions on the full data set from the trace conditioning rats (**Figs. 3B, 4E**). Both exponential equations performed better than a linear function based on the full data set, with the four parameter Exponential 2 equation providing a significant improvement relative to the linear function (**Fig. 4F**). Taken together, these data coupled with prior reports illustrate how exponential functions best account for the relationship between dopamine and behavioral responding on the order of 10s of seconds.

## Discussion

We demonstrate that the CS dopamine response in the NAc encodes the trace period between the end of the cue and the delivery of the reward. These results are consistent with the findings from Pavlovian delay conditioning and operant delay discounting tasks where the CS dopamine response signals an estimate of reward value. Specifically, these studies demonstrate that the CS-evoked *increase* in dopamine signaling is inversely related to the time until the reward (Fiorillo et al., 2003; Roesch et al., 2007; Kobayashi and Schultz, 2008; Day et al., 2010; Gan et al., 2010). However, we find that a reward predictive cue produces a *decrease* in NAc dopamine levels when it signals a long trace period. This cue-evoked decrease in dopamine signaling is unexpected, as the presentation of a CS-during delay conditioning can still enhance dopamine release due to response generalization (Day et al., 2007). Furthermore, a recent study found no change in dopamine neuron activity to cues that are reliably delivered after the reward delivery in animals well-trained on a backward conditioning task (Taira et al., 2024). Based on these findings from delay and backward conditioning tasks, one would expect either no dopamine response or a modest increase in dopamine levels to a cue that is never paired with a reward delivery. However, by examining the dopamine response during a trace conditioning task, we uncovered that dopamine encoding of reward value is not solely limited to positive changes in dopamine levels, but rather can encompass both positive and negative changes in the dopamine response to the CS.

Animals reliably prefer cues associated with immediate rewards relative to cues associated with delayed rewards in operant tasks (Roesch et al., 2007; Day et al., 2010; Gan et al., 2010). In agreement, rats exhibited a higher level of conditioned responding on Short Trace trials relative to Long Trace trials. We note that the magnitude of conditioned responding during trace conditioning was ~5 times lower relative to similarly designed delay conditioning tasks that we have used previously, which indicates that the temporal separation between the CS and US during Pavlovian conditioning influences the intensity of responding during the cue (Fonzi et al., 2017; Stelly et al., 2020; Stelly et al., 2021; Lefner et al., 2022a; Lefner et al., 2022b). The difference in conditioned responding between trial types emerged after ~10 training sessions and was driven by an increase in responding to the Short Trace CS. In contrast, a difference in the latency to respond between trial types rapidly emerged during early training sessions and was driven by a slowing in the response to Long Trace CS. These data underscore how distinct cue-evoked behavioral responses can develop independently of one another during Pavlovian conditioning. In support, we previously found that systemically antagonizing dopamine receptors can differentially regulate conditioned responding and the latency to respond during delay conditioning (Lefner et al., 2022b). Future studies are needed to determine how cue-driven behaviors during trace conditioning are controlled by dopamine transmission.

Short Trace trials produced higher levels of conditioned responding and greater CS-evoked dopamine release in the NAc relative to Long Trace trials. However, we found no correlation between conditioned responding and the CS dopamine response during trace conditioning. These results are consistent with our prior work using well-trained animals undergoing delay conditioning (Fonzi et al., 2017). In this study distinct cues signaled the time since the previous reward delivery. One audio cue always followed a short ITI and signaled a higher rate of reward while another audio cue always followed a long ITI and signaled a lower rate of reward. While there was greater dopamine release to the short ITI cue, this was not accompanied by a higher level of conditioned responding in well-trained subjects under static conditions. Rather, CS-evoked dopamine release in this delay conditioning task correlated with conditioned responding during early learning and when the training context was changed (Fonzi et al., 2017; Stelly et al., 2021). Taken together, the absence of a relationship between NAc dopamine levels and conditioned responding is to be expected given that dopamine recordings during this trace conditioning task were performed in well-trained animals under static conditions.

The dopamine neuron response to reward predictive cues adjusts accordingly to the range of options that are presented (Tobler et al., 2005). For example, a cue signaling a medium-sized reward will evoke a large dopamine response when an alternative option is a small-sized reward. However, the same cue signaling a medium-sized reward will evoke a small dopamine response when an alternative option is a large-sized reward (Tobler et al., 2005). Our previous research illustrates that cue-evoked dopamine release in the NAc is similarly influenced by the range of options that are presented (Fonzi et al., 2017). In this study, rats were trained on a delay conditioning task where distinct cues were associated with different reward rates. CS-evoked dopamine release in the NAc encoded the relative reward rate when the different trial types were experienced in either the same session (context-dependent) or in separate sessions (context-independent). However, the overall magnitude of CS-evoked dopamine release was smaller in rats that were trained to experience the trials in separate sessions (Fonzi et al., 2017). Taken together, these findings suggest that the context in which reward-predictive cues are presented can influence the *magnitude* of the dopamine response to that cue, but the *direction* of dopamine response is context-independent. Given that rats experienced both Short Trace and Long Trace trials within the same session, it is likely that a within-session contrast effect between trial types influenced the magnitude of the dopamine signals, but not the direction of the observed dopamine responses.

Whereas there was no link between dopamine and conditioned responding, we instead found that CS dopamine levels in the NAc were related to the response latency. Our initial analysis illustrated a significant linear correlation between dopamine levels and the latency. However prior research indicates that the relationship between dopamine and behavior over longer time scales may be better explained by an exponential function (Fiorillo et al., 2003; Kobayashi and Schultz, 2008). Indeed, we found that best fit exponential functions were more accurate at predicting dopamine levels relative to a standard linear function in well-trained animals. Future studies are needed to determine how CS-evoked dopamine relates to the response latency during early training sessions.

A regrettable limitation of this current study was the use of only male subjects, though we note that the data were collected prior to the implementation of the sex as a biological variable policy from the NIH. Sex differences have been observed in a number of dopamine-dependent behaviors (Lynch and Carroll, 1999; Zachry et al., 2019; Kutlu et al., 2020; Bishnoi et al., 2021; Chen et al., 2021; George et al., 2021). In particular, females exhibit higher levels of conditioned responding and a faster response latency than males during a delay conditioning task (Lefner et al., 2022a). Increasing evidence illustrates how cycling hormones can modulate dopamine transmission, which can subsequently impact dopamine-dependent behaviors (Calipari et al., 2017; Yoest et al., 2019; Zachry et al., 2021; Zhang et al., 2021). Therefore, future research is needed to determine how NAc dopamine signaling relates to behavioral responding during trace conditioning in females across the estrus cycle.

Prior research highlights that midbrain dopamine neurons contribute to evaluating differences in time on rapid timescales (< 2s) (Pasquereau and Turner, 2015; Soares et al., 2016). For example, dopamine neuron activity can reflect the expectation that an event will occur (Pasquereau and Turner, 2015). Additionally, optogenetic manipulations of dopamine neurons can bidirectionally alter the perceived temporal interval between two cues (Soares et al., 2016). Our current results extend these findings to illustrate that the CS-evoked dopamine response in NAc core can encode the trace period over long temporal intervals (0-55 s). Collectively, these highlight a role for dopamine in timing-related calculations over a range of timescales.

