## Supplemental Materials for "Nucleus accumbens dopamine encodes the trace period during appetitive Pavlovian conditioning"

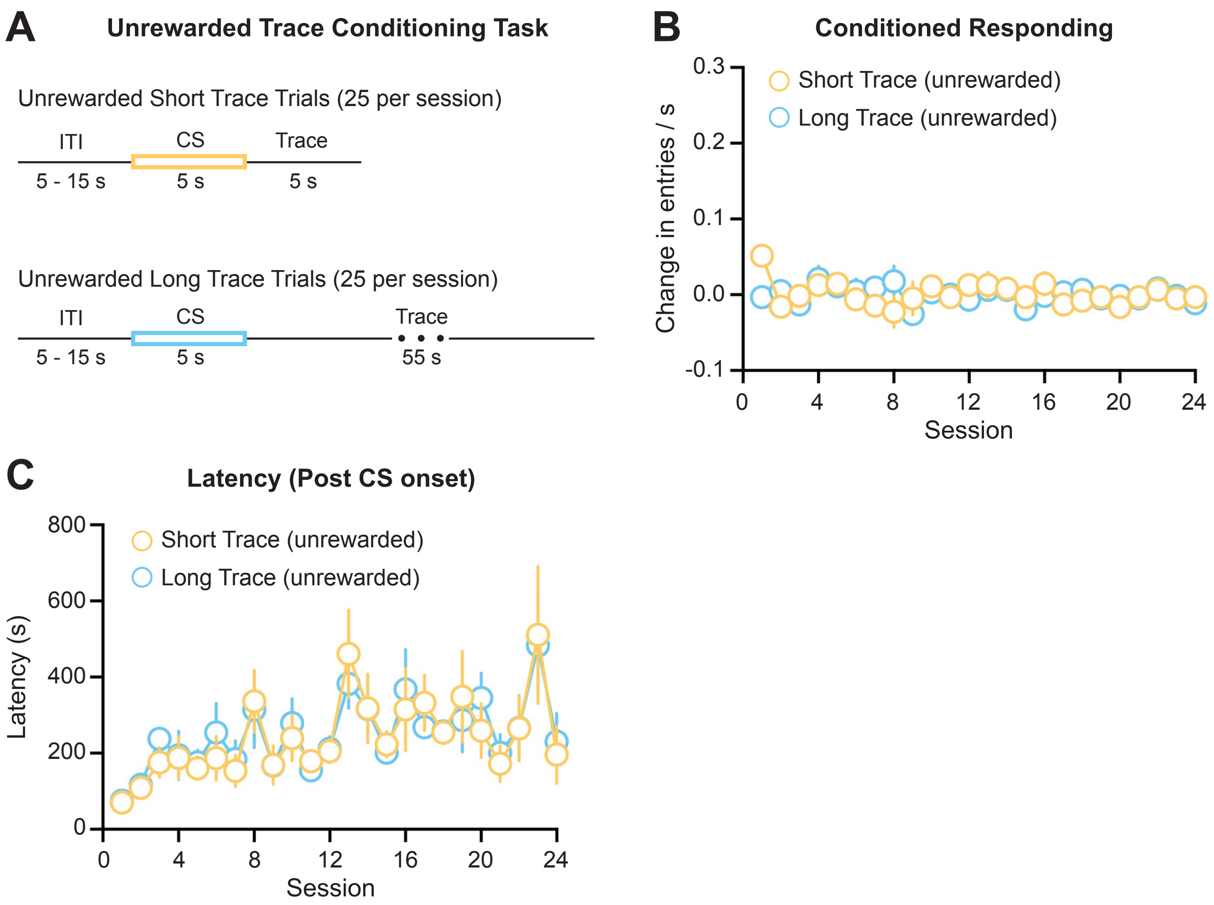


**Extended Figure 1-1.** Unrewarded trace conditioning task. (A) Schematic of the Unrewarded Short and Long Trace trials, which are presented in a pseudorandom pattern within a session. (B) No change in conditioned responding to unrewarded cues across training sessions (N = 5 rats; 2 male and 3 female). (C) Response latency following the CS presentation across training sessions.

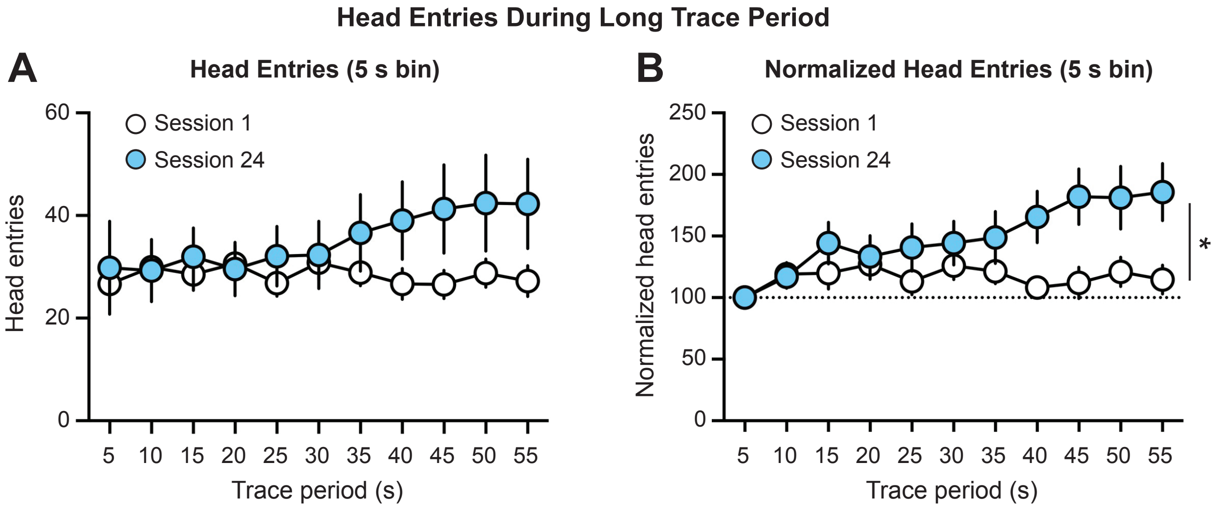


**Extended Figure 1-2.** Head entries during the long trace period. (A) Head entries increase throughout the trace period over training (two-way ANOVA: interaction effect $F_{(10,220)} = 4.4$, $p < 0.001$). (B) Head entry data normalized to the responding during the first 5 s of the trace period illustrates an increase in responding throughout the trace period over training (two-way ANOVA: interaction effect $F_{(10,220)} = 4.6$, $p < 0.001$; training effect $F_{(1,22)} = 6.6$, * $p = 0.02$).

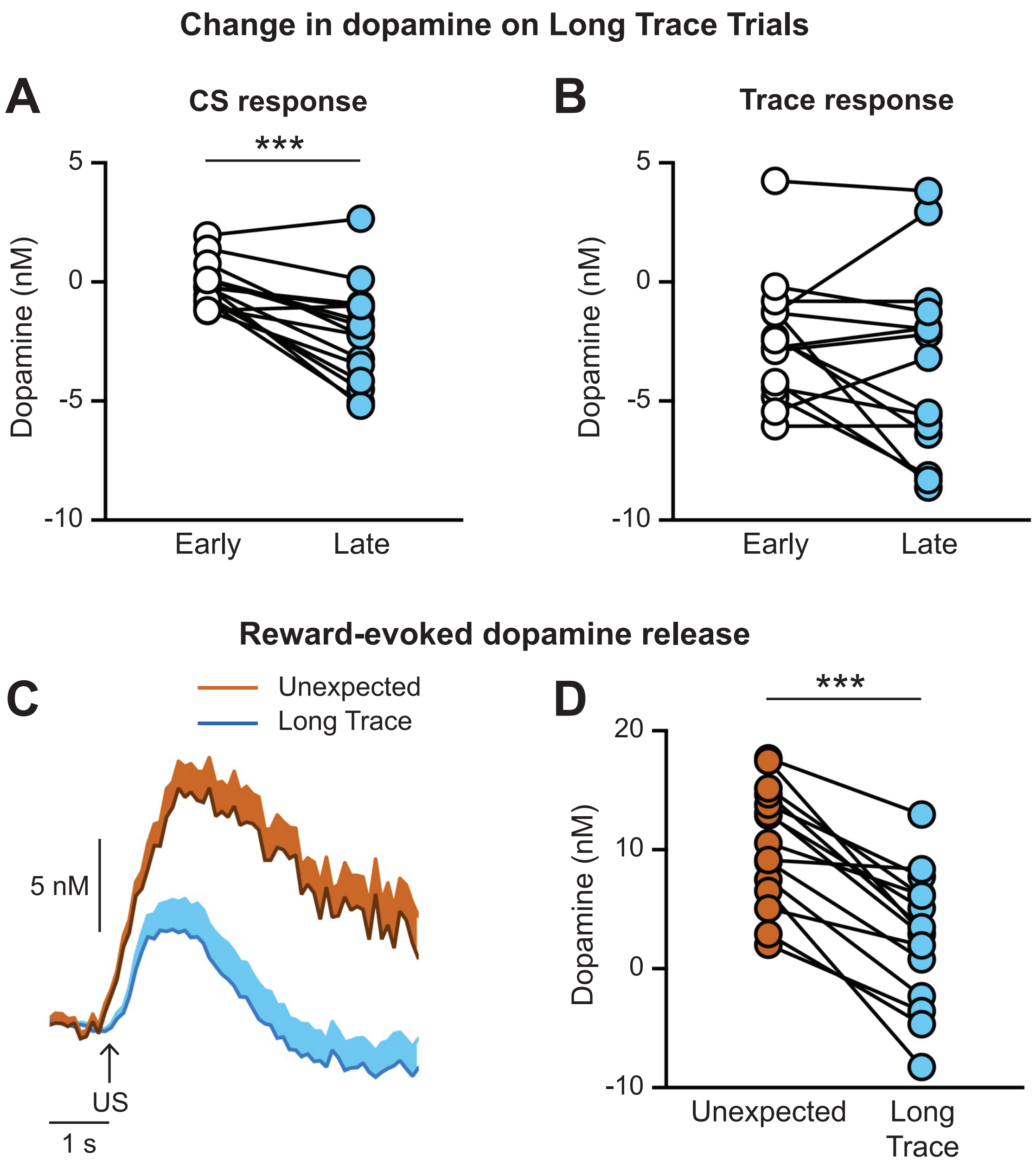


**Extended Figure 2-1.** Additional analyses of NAc dopamine signals. (A,B) Change in dopamine on Long Trace trials during the CS period (A) and the first 5 s of the trace period (B). Values were calculated by examining the average dopamine signal during the first 1 s and last 1 s of the epoch. Dopamine levels significantly decreased during the CS presentation (A, paired t-test: $t_{14} = 4.9$, *** $p = 0.0002$), but had no further change during the initial trace period (B, paired t-test: $t_{14} = 1.5$, $p = 0.14$). (C,D) US-evoked dopamine release on Long Trace trials is significantly smaller than the dopamine response to an unexpected food pelleted delivered outside of the Pavlovian conditioning sessions (paired t-test: $t_{14} = 7.7$, *** $p < 0.0001$).
